# Viscous effects and complex local flow behaviors dominate hemolymph circulation in the living wings of locusts

**DOI:** 10.1101/2021.09.15.460448

**Authors:** Mary K. Salcedo, Brian Jun, Pavlos Vlachos, Lakshminarayanan Mahadevan, Stacey A. Combes

## Abstract

An insects’ living systems – circulation, respiration, and a branching nervous system – extend from the body into the wing.^1,2^. Hemolypmh circulation in the wing is critical for hydrating tissues, such as the highly elastic resilin^3^ that enhances wing flexibility, and for supplying nutrients to living systems, including sensory organs such as scent-producing patches, sound-receiving tympana, and wind-sensing sensilla distributed across the wing.^4–7^During flight, the presence of hemolymph in the wings reduces aerodynamic instabilities like flutter^8,9^, and faster hemolymph flows are induced by flapping.^10^ Despite the critical role of hemolymph circulation in maintaining healthy wing function, wings are often considered “lifeless” cuticle, and most measurements remain qualitative or employ coarse, bulk-flow techniques. While pioneering work in the 1960s mapped hemolymph flow direction in 100 insect species,^11^ half a century later we still only have quantitative measurements of flow within the wings of a few insects. Here, we focused on the North American locust *Schistocerca americana*, a well-studied agricultural pest species, and performed a detailed, quantitative study of global and local hemolymph flows in the densely venated fore and hind wings, along with key regions in the body and pumping organs. Through high-speed fluorescent microscopy, we measured 800 individual trajectories of neutrally buoyant fluorescent particles that move in sync with hemolymph, in the wings and body of 8 live, resting locusts. Our data show that overall flow within the wings is circuitous, but local flow behavior is highly complex, with three distinct types of flow (pulsatile, continuous, and “leaky”) occurring in various combinations in different areas of the wing. We provide the first quantitative measurements of “leaky” flow into wing regions that act as sinuses, where hemolymph flows out of tubular veins and pools within thin membranous regions. We also calculate Péclet, Reynolds, and Womersley numbers, and find that viscous effects dominate flow regimes throughout the wing. Pumping organs and wing regions closest to the body display significantly faster flows and higher Reynolds numbers, but remain within the viscous flow regime. Given the central role of wings in sustaining ecologically important insect behaviors such as pollination, migration, and mating, along with the vast diversity of insect wings seen in nature, this first detailed, quantitative map of hemolymph flows across a wing provides a template for future studies investigating the dynamics of hemolymph flows critical to sustaining wing health among insects.

## Introduction

Functioning and healthy insect wings are inextricably linked to active hemolymph circulation within the wings.^1, 2, 7^ If one were to cut an imaginary slice through an insect wing, this would show that an insect’s circulatory, respiratory, and nervous systems all extend from the body into the wing veins (Fig. 1). Hemolymph serves to hydrate tissues, supply the nervous and respiratory systems, and circulate cells involved in immune function, and hemolymph flow is a critical hydraulic tool during insect growth, metamorphosis, and wing expansion.^12^ Recent work has also confirmed that hemolymph circulation is necessary for the scent-producing organs on lepidopteran wings^4^ and the hundreds of sensory hairs distributed across dragonfly wings to function properly.^6, 10^

**Figure 1.**
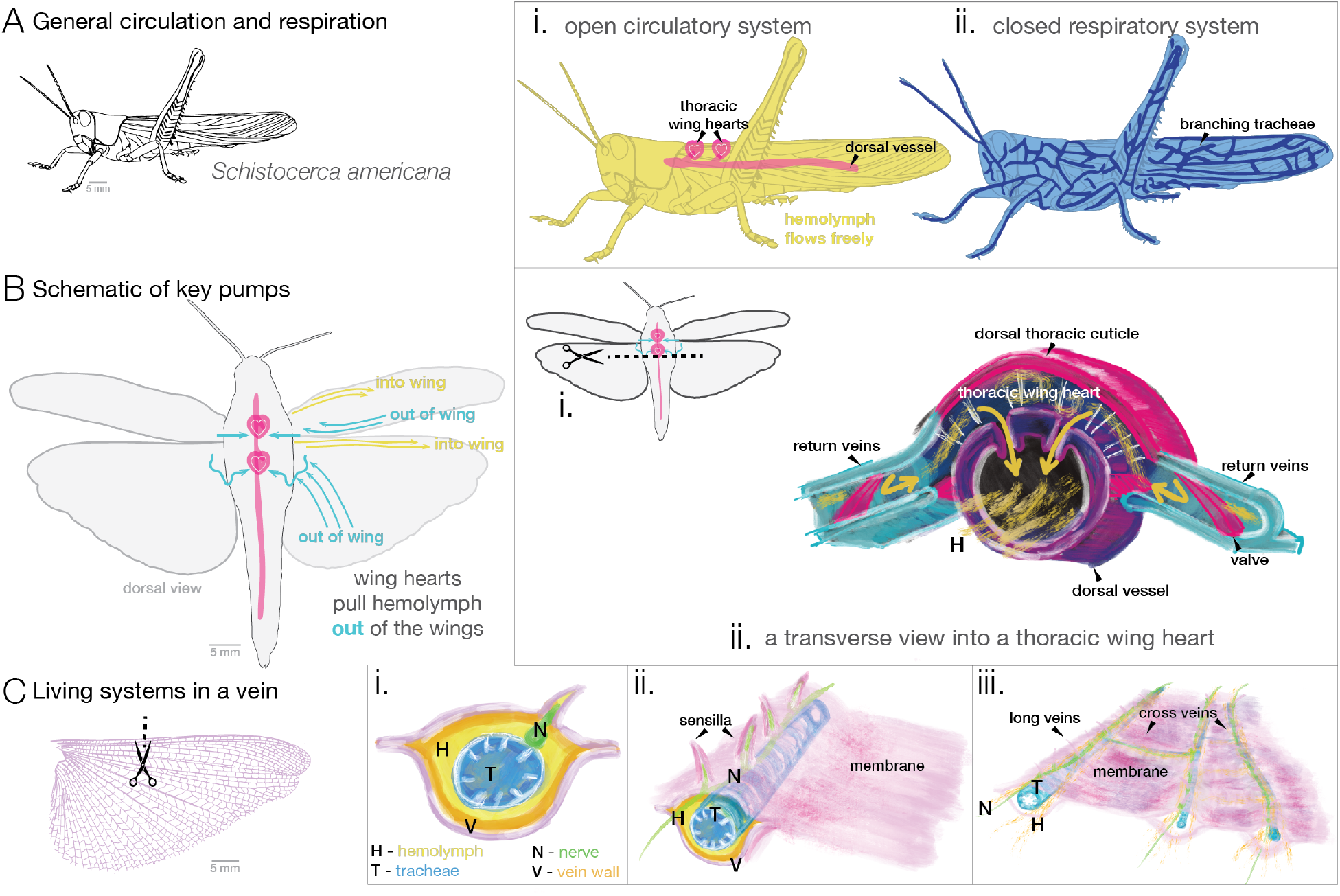
Physiology and fluid systems in the grasshopper *Schistocerca americana* and its wings. **(A)** The two main fluid systems within an insect include its open circulatory system (i) and its closed respiratory system (ii). Within an open circulatory system hemolymph (insect blood) is from pumped posterior to anterior via a long tubular heart called the dorsal vessel. Accessory hearts (i.e., pumps) in the thorax called “wing hearts” pump blood from the wing.^18^ An insect’s respiratory system is network of tracheal tubes which connect directly to tissues transporting oxygen and carbon dioxide by advection and diffusion.^19^ **(B)** In *S. Americana*, thoracic wing hearts have “return conduits” (i.e., scutellar branches) where hemolymph leaves the wing and returns to the main heart. A transverse slice through the thorax (i) reveals how these thoracic wing hearts (ii) are located dorsally above the main tubular heart. **(C)** Take an imaginary slice through a vein and it reveals (i) hemolymph, tracheal branches, nerves, and vein wall.^1^ Extended views (ii-iii) shows nerve branches connecting to proprioceptors, and how hemolymph and tracheal tubes form networks inside the wing. Inspired by Pass.^1, 2^

Despite this wide range of important physiological functions, insect wings have primarily been studied in relation to their mechanical function during flight. Structurally, insect wings are composed of chitinous, tubular veins and thin, membranous regions^13^. The mechanical behavior of insect wings is influenced not only by the wing’s material properties and the pattern of supporting wing veins, but also by the presence, and perhaps movement, of hemolymph within the veins. Recent studies have shown that approximately 30% of an insect’s total hemolymph will cycle through the wings^14^, and that hemolymph circulation is enhanced by flapping, with hemolymph moving more quickly through flapping dragonfly wings than still wings.^10^ This study also showed that large wing structures, like the dragonfly’s pterostigma, hold copious amounts of hemolymph, but the mechanism of hemolymph flow into these structures has not been examined. In dynamic 3D models of lady beetle wings, the presence of hemolymph was shown to reduce aerodynamic instabilities like flutter.^9^. Wing flexibility plays a critical role in the mechanics of flapping wings, and flexibility is correlated with high concentrations of resilin, a hyper-elastic protein that requires hydration to maintain its elasticity.^3^ Some insect wings, depending on venation pattern, morph and change shape to unfold prior to or during flight.^15^ Beetles can initiate wing unfolding with a pulse of hemolymph,^16^ a mechanism that may also be used by earwigs to unfurl their fan-shaped, resilin-rich wings.^17^ While the mechanical properties of insect wings are relatively well-studied, the internal, living systems within wings (Fig.1A) have been largely ignored, despite their critical importance for maintaining wing mechanical properties and physiological functions.

An insect’s open circulatory system (Fig. 1A,i) hydrates tissues and delivers nutrients throughout the body and appendages (legs, antennae, and wings), with flow driven primarily by a long, tubular pump (i.e., heart) known as the dorsal vessel, which pulls fluid from the posterior end of the open body cavity and pumps it into the anterior end.^12^ Hemolymph contains hemocytes that are involved in immune functions such as phagocytosis, encapsulation, and clotting in response to damage^20^, but hemolymph does not play a major role in gas exchange (as blood does in vertebrates), despite the presence of the respiratory pigment hemocyanin in many insects^21^. Instead, the tracheal system (Fig. 1A,ii), a branching network of tubes, delivers oxygen directly to tissues throughout the body and appendages via diffusion and advection (bulk flow).^19^ Tracheal branches (an insect’s respiratory network) first extend into wing tissue during wing pad development and can be found in most adult wing veins.^22^ In addition to the dorsal vessel that pumps from the posterior to anterior body cavity, insects have smaller thoracic “wing hearts” (Fig. 1B), which are pulsatile organs (i.e., pumps) that pull hemolymph out of the wings, driving overall circulation in the wings that can be either circuitous (e.g., entering through the leading edge and returning to the body through the trailing edge (Fig. 1B).^7, 11, 23^) or tidal (alternately entering and then being pulled out from all of the veins in distinct pulses).

In 1960, John Arnold examined the wings of 100 insect species and described the presence and overall patterns of hemolymph circulatory flow in insect wings^11^ This in-depth study and review was an attempt to resolve 200 years of arguments concerning whether or not wing circulation existed. Since this seminal study, only a handful of studies have examined the importance of hemolymph circulation to the mechanical or physiological processes of the wing. Some trends concerning hemolymph circulation are well understood; Arnold described two main flow patterns across 14 insect orders, in at-rest insects: circuitous flow (circuit-like) and tidal (in all veins at once, and then out). Further studies by Wasserthal detailed bulk tidal flow in resting lepidoptera, describing how large moths like *Attacus atlas* (with a wing span of 30 cm) use accessory pumps (thoracic wing hearts), thoracic air sacs, and tracheae extending into the veins to push hemolymph into all wing veins.^24^ Chintapalli and Hillyer used fluorescent beads to study hemolyph flow in mosquito wings (*Anopheles gambiae*, with a wing span of only 1 mm), describing the distinctive, pulsating and circuitous flow route within these tiny wings, which is driven by an independent thoracic wing heart (i.e., an extra pump).^23^ Wang et al., the first study to measure bulk hemolymph flow in flapping wings, showed that flapping induces faster hemolymph flows than those observed during rest.^10^ These three works have established clear connections between an insect’s pumping systems, air network, and the circulation of hemolymph in at-rest and flapping wings.

However, there is still a significant lack of quantitative analyses of hemolymph circulation within wings, particularly concerning local flow behaviors within the veins, as opposed to global, bulk flow throughout the entire wing. Many past studies have focused on bulk-flow measurements in insects at rest^10, 16, 25^, as measuring fluid movement within tiny insect wing veins is not a trivial task^7^. However, the use of injected fluorescent particles to track fluid flow has increased over the last decade^7, 23, 26, 27^, and we hypothesized that with the appropriate choice of particle size and buoyancy, this technique could be used to produce a comprehensive hemodynamic map of wing circulation that accounts for local as well as global flow behaviors.

Here, we describe the results of a study in which we used high-speed, fluorescent microscopy to observe and quantify active hemolymph circulation within the densely venated fore- and hindwings, as well as the body of live, adult locusts *Schistocerca americana* at rest (Fig. 2). We chose this species because its has commonly been used as a model organism for studies of flight; *S. americana* is known for its swarming behavior and its genus has been well-studied in terms of wing biomechanics.^28–30^ *Schistocerca* employ an “umbrella-effect” during flight, in which the forewings and hindwings flap in anti-phase and the corrugated hindwings balloon out, flexibly deforming with each wing flap.^31, 32^ In addition, this species is of intermediate size (in relation to previous wing circulation studies), and is an ecologically relevant pest species, making it readily available through collaborations with USDA (U.S. Department of Agriculture) facilities.

**Figure 2.**
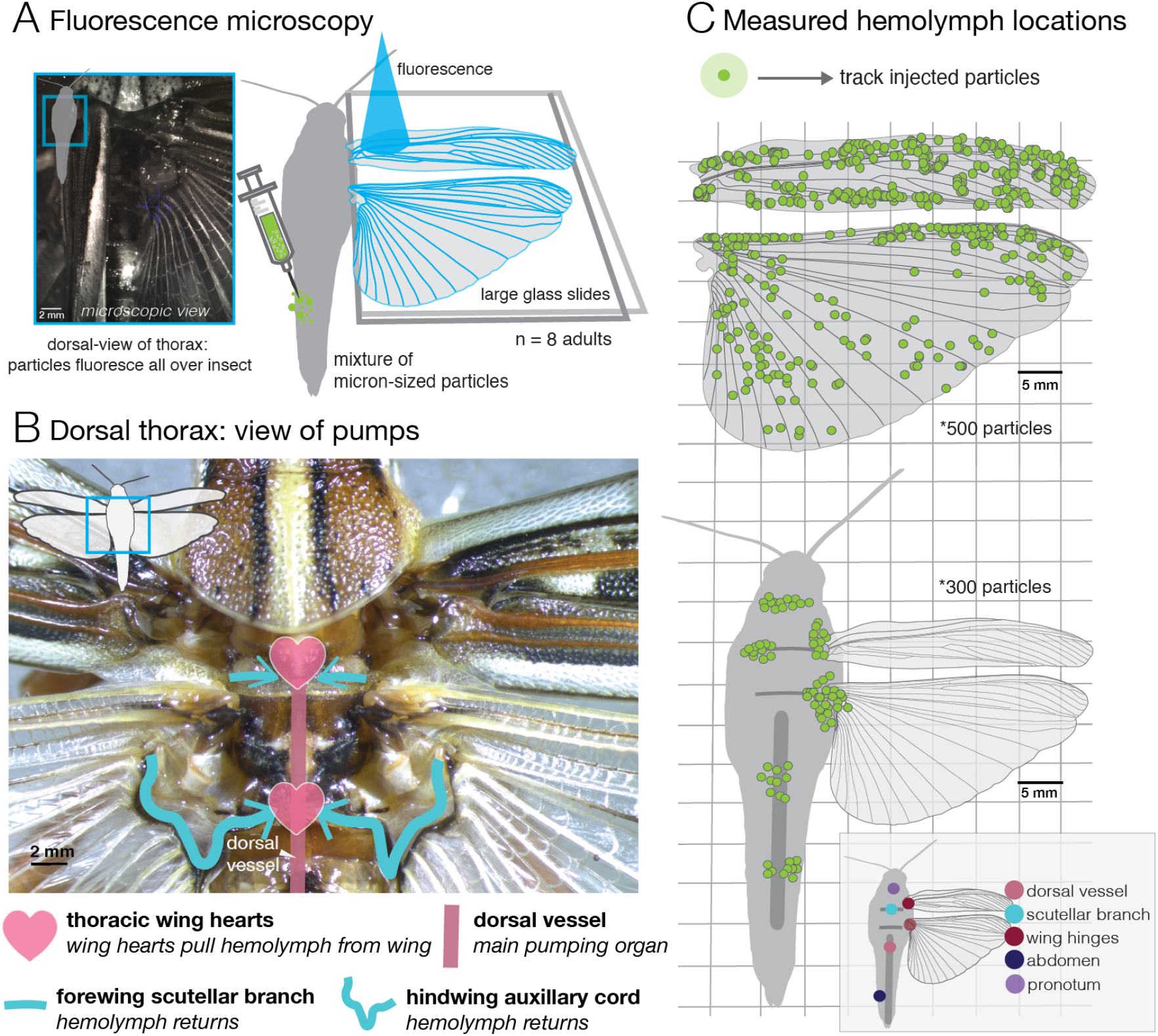
Tracking hemolymph via fluorescent particles in *S. americana*. **(A)** View of dorsal thorax of grasshopper under a fluorescent microscope (left). Insects were injected with neutrally buoyant fluorescent particles. Before imaging and particle injection, *S. Americana* are briefly anesthetized with carbon dioxide and quickly restrained with modeling clay; wings were spread between two glass slides (blue light is fluorescence). **(B)** Dorsal view to indicate location of thoracic wing hearts and return conduits into the wing heart pump. The dorsal vessel dominates pumping hemolymph within an insect but cannot circulate hemolymph into the wings without assistance of thoracic wing hearts. **(C)** Top map (normalized coordinates) of all particles (500 total) tracked and quantified across 8 adult grasshoppers in both the forewing and hindwing (16 wings total). Bottom map of measured particles (300 total) across the body.

We investigated four key questions: **(1)** What is the overall pattern of flow for hemolymph moving into and out of the wing? **(2)** What local flow behaviors are present in the wing at smaller scales, and how does flow behavior vary regionally? **(3)** How does peak flow velocity and pulsing rate vary across the body and wings? **(4)** How do the fluid dynamic regimes of flow vary across the wing?

## Results

### Tracking hemolymph in every vein and local patterns

We successfully adapted fluorescent microscopy techniques^23^ to construct a comprehensive hemodynamic analysis of circulation in the wings and wing pumps of *S. americana* (Fig. 2A). We injected and tracked neutrally buoyant particles moving with hemolymph throughout the major regions of the wings and bodies (Fig. 2B) of 8 living, at-rest *S. americana* adults, producing a total of 800 particle trajectories (Fig. 2C). The forewing of locusts, the tegmen, is a thickened, semi-leathery wing that covers the larger hindwing ( 2.5x greater area), which is folded like a corrugated fan under the forewing when the insect is at rest. Both wings are densely venated, and contain longitudinal veins spanning the wing from base to tip, which are interconnected by numerous, shorter cross-veins. Particularly near the wing base, wing veins are not circular in cross-section, but rather u-shaped and shallow (Supplementary Movie 1). Based on structural similarities between the wings, we identified five distinct wing regions, within which we assessed local flow characteristics. These regions include the 1) leading edge (largest diameter veins), 2) membrane (a large sinus present between wing layers near the leading edge), 3) wing tip (small diameter, highly interconnected veins), 4) lattice (mostly orthogonally connected veins), and 5) trailing edge (larger diameter veins) (Fig. 3B).

**Figure 3.**
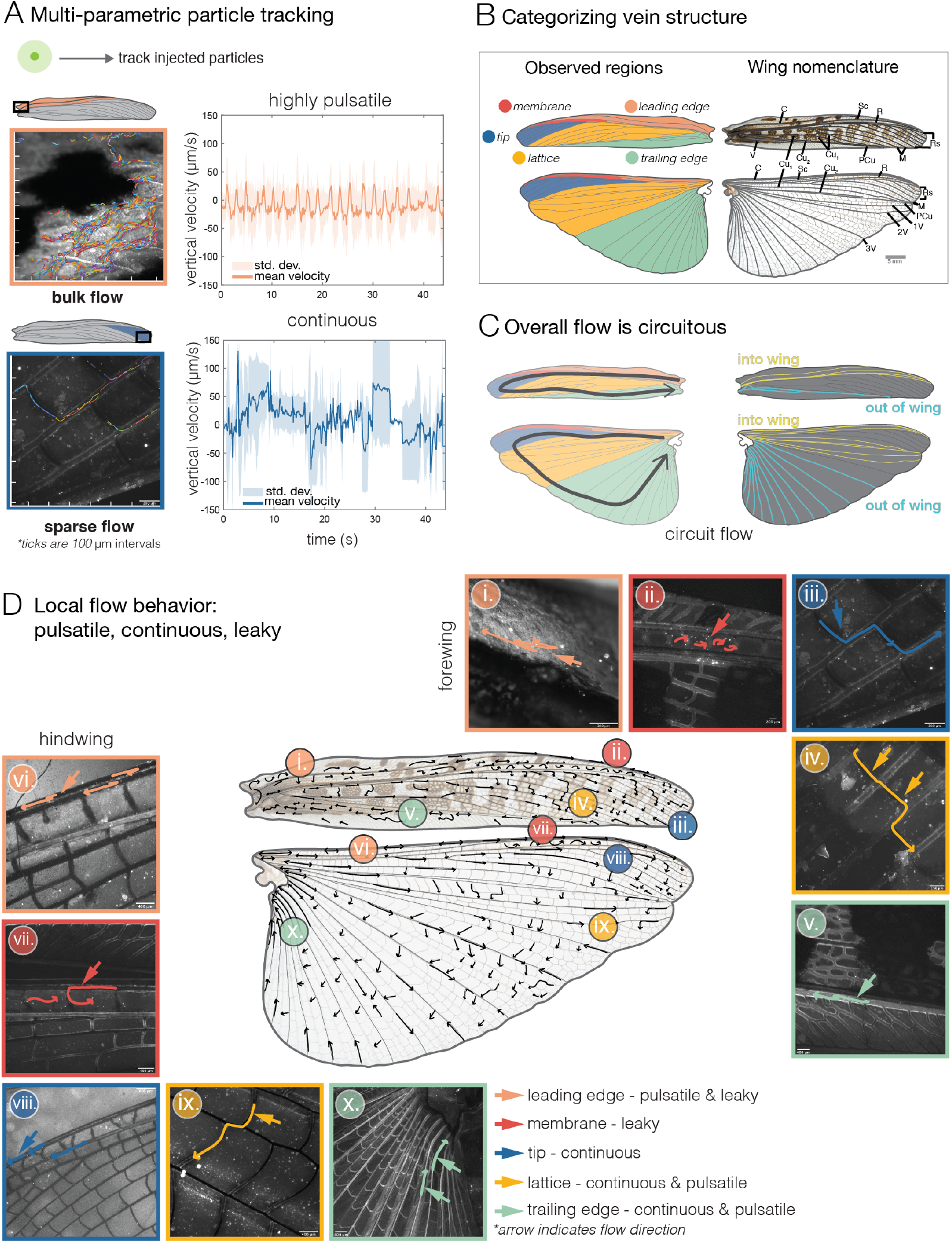
Circuitous flow pattern and local flow behavior. **(A)** Using multi-parametric tracking of cells to detect bulk movement of particles allows for tracking of large numbers of particles. Flow near the wing base (top) shows distinct pulsatility while in regions like at the wing tip (bottom), flow is traversing more junctions, and patterns are not as clear. **(B)** To categorize vein structure wing metrics were simplified into five regions (left) based on vein location and structure: 1) leading edge (pink, costa to subcosta), 2) membrane (red, subcosta to radius), 3) wing tip (dark blue, radial sector to medius), 4), lattice (yellow, medius to post cubitus), and 5) trailing edge (light green, post cubitus to vannal region). Labels follow long vein nomenclature (short veins are typically unnamed).**(C)** Overall flow in the wing was found to be circuitous where hemolymph moved into the wing through C, Sc, and R veins, and out of the wing via the Cu and V veins. **(D)** Following Arnold’s (1964)^11^ wing drawings, hand-drawn vectors represent hemolymph behavior (based on tracking analysis). Tracking fluorescent particles reveals that flow behaves in three modes: pulsatile (double-headed arrow), leaky (curved arrow), and continuous (straight arrow). Examples of forewing venation (i.-v.) and hindwing (vi. - x.) in each of the five regions. *Wing veins: C - costa, Sc - subcosta, R - radius, Rs - radius sector, M - medius, Cu - cubitus, PCu - post cubitus, V - vannal*

Using fluorescent microscopy and high-speed video, we recorded fluorescent particles flowing in sync with hemolymph that was advected into the wing at the wing base (Supp. Movie 1), and we employed multiple tracking methods (Fig. 3A), including multi-parametric particle tracking and semi-automated tracking,^33^ to determine flow velocities and patterns. We found that in *S. americana*, hemolymph is pumped out of the forewing and hindwing near the trailing edge by each wing’s respective thoracic wing heart (Fig. 2B), and hemolymph flows passively into the wing from the thoracic space, entering through the largest, leading edge veins, the costa, subcosta, and radius. This creates an overall circuitous bulk flow pattern in both the fore- and hind wings 3C.

### Local flows are pulsatile, continuous, and leaky

Although bulk flow within locust wings can be described as a one-way circuit, with hemolymph entering via leading edge veins and exiting from trailing edge veins (Fig. 3C, Supp. Movies 1 and 6), local flow behaviors within the veins are complex and time-varying. Hemolymph does not travel along simple, predetermined paths through the wing, but rather may display one of several local flow behaviors at any given vein junction, at a particular point in time. Specifically, while measuring and tracking active hemolymph circulation in every vein within both the forewing and hindwing, we observed three distinct, local flow behaviors: pulsatile, continuous, or leaky flow (Supp. Movies 1-6). Multiple types of flow behaviors can be found within many of the wing regions, and the occurrence of some local flow behaviors appears to be a function of proximity to the insect body and associated pumping organs.

Flow is pulsatile in much of the wing (Supp. Movies 2 and 6), with particles pulsing forward and then stopping or reversing direction for a shorter distance, at regular intervals. As a result, hemolymph can move in two or more alternative directions at many vein junctions, and flow can appear tidal in some smaller veins (Supp. Movie 5). Generally in insect wings, leading edge veins are large and tend to decrease in diameter across the span and chord (i.e., along the length and width of the wing). We found that hemolymph movement is dominated by pulsatility in the leading edge, where the veins are large in diameter (170-250 *μ*m). In sync with the wing hearts (0.9 Hz), hemolymph flow in the leading edge region pulses back and forth rapidly within a vein (Supp. Movie 8), with net movement eventually proceeding towards the wing tip and downward through cross veins.

Leaky flow, a unique flow behavior in insect wings that we quantify here for the first time, occurs when particles move out of the wing veins and into an adjacent membranous region (a large sinus region), eventually flowing back into veins from the sinus (Supp. Movies 2 and 3). What we characterized as the “membrane” region of the wing occurs at approximately two-thirds of the wing span, towards the leading edge, and in this region hemolymph flows out of the leading edge veins (costa, subcosta, and radius) and into the pocket-like membranous sinus (Supp. Movie 6). Particles moving from vein to membrane in this region maintained velocities similar to those in the tubular veins of the leading edge. This “pseudo-stigma”^34^ (similar to a dragonfly’s pterostigma) is a significant feature in both the forewings and hindwings of *S. Americana* (Fig. 3D ii and vii, membrane region). Within the leading edge and membrane regions of the wing, we documented both pulsatile and leaky flow behaviors (Fig. 3D, i/vi and ii/vii, Supp. Movies 1 and 2).

Continuous flow occurs where particles move in one direction continuously; velocity may increase and decrease in sync with hemolymph pulsing, but the particles never stop entirely or change direction. We observed continuous flow behavior, as well as pulsatile flow, within the remaining three wing regions - the wing tip, lattice, and trailing edge regions (Supp. Movies 4-6) (Fig. 3D, iii/viii, iv/ix, and v/x). Pulsation tends to be damped within the wing tip and lattice regions, with flow more often moving continuously towards the trailing edge, whereas pulsatile flow is more common within the trailing edge region, where hemolymph is pumped out of the wing.

### Wing hemolymph moves faster in regions near the body

Hemolymph within the wing flows faster in regions near the body and slower in regions toward the tip of the wing. We calculated peak and median particle velocities to compare how quickly hemolymph moves in different wing regions. (Fig.4). Overall, flow velocities are higher in the hindwing as a whole (Fig.4A), likely due to its larger size. Within each wing, the highest peak flow velocities occur in regions near the wing base; in the forewing, the highest peak velocities occur in the trailing edge region (where the thoracic wing hearts pull hemolymph out of the wing through the scutellar branch) and in the hindwing, the highest peak velocities occur in the leading edge region, where hemolymph is drawn into the wing from the thoracic cavity to replace fluid pumped out at the trailing edge. Median flow velocities show similar patterns (Fig.4B), but with smaller differences between regions.

**Figure 4.**
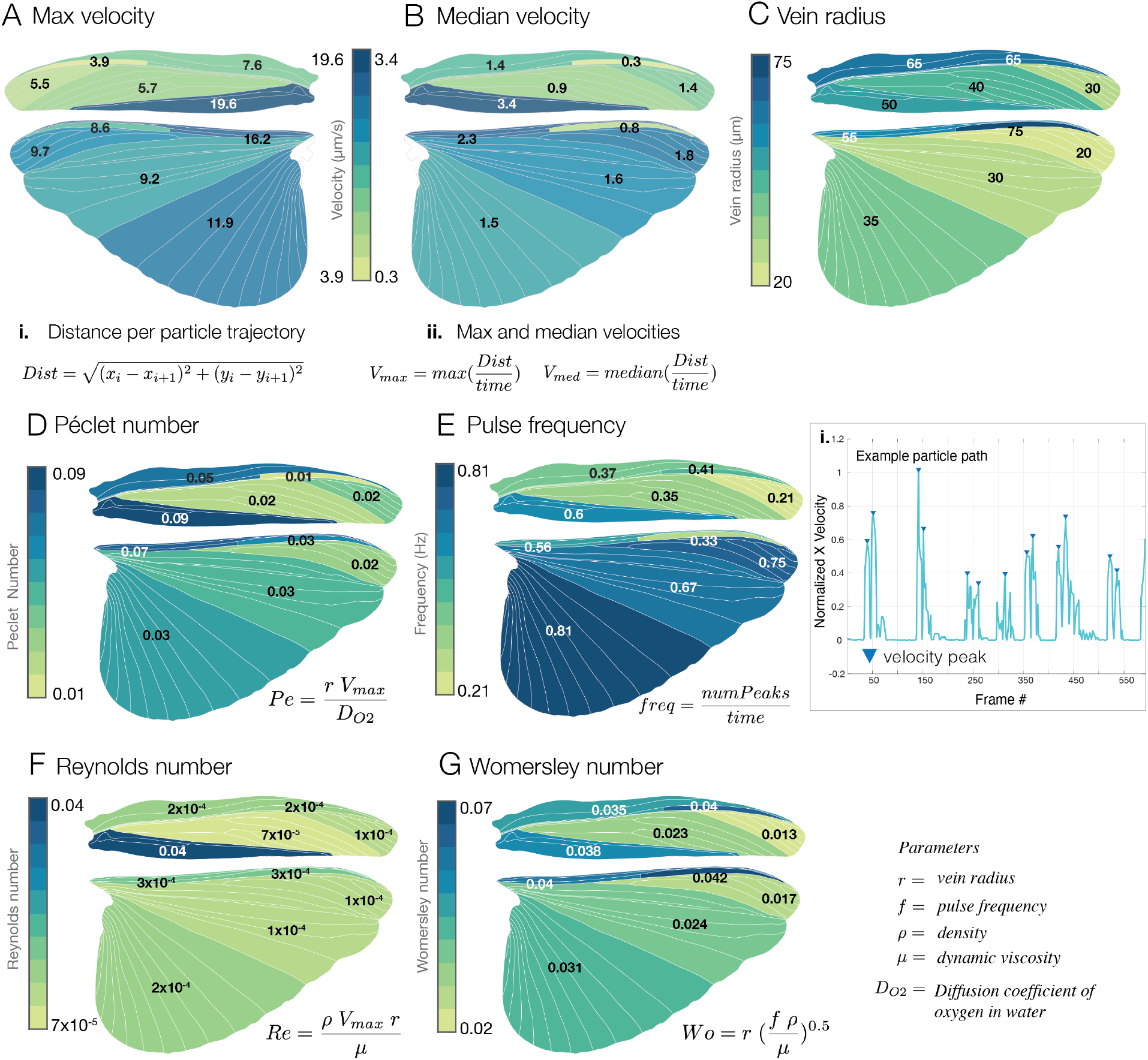
Flow dynamics across wing regions. **(A)** Maximum velocities per wing region (i, ii). Faster flows occur in the trailing edge of the forewing and leading edge of the hindwing. **(B)** Median velocities per wing region. **(C)** Average vein radius per region (n = 25 vein radii measured and average taken). **(D)** Péclet number for each region where the diffusion coeffcient is for oxygen in water. **(E)** Pulse frequency is calculated by the number of velocity peaks over time (i). **(F)** Reynolds number describes a viscous flow regime in all regions. **(G)** Womersley number across regions mirrors Womersley flow in arterioles/venules.^35^ *Medians of the wing regions are represented in A,B,D-G. Per wing figure - 8 individual insects and 500 digitized particles*.

Structurally, veins in the leading and trailing edge regions of the forewing are wider than those in the forewing tip and membrane regions. At the wing tip, flow follows the perimeter vein and also begins moving down the chord of the wing into the lattice and trailing edge regions. In both the fore- and hindwings, hemolymph flow slows significantly in the wing tip region (Fig.4A,B) and increases again in the trailing edge region (paired t-test, P<0.05). In the hindwing, the anal veins within the trailing edge region serve as long conduits (with fewer junctions to traverse) that all feed into the same return conduit (i.e., auxillary cord), where flow is pulled out by the posterior thoracic wing heart (Fig.2B).

### Viscous effects dominate flow regimes in the wings

Based on our measurements of flow velocity and vein size, we found that hemolymph flow is dominated by viscous effects within all of the wing veins of *Schistocerca americana*, despite the relatively large size of this insect. In Fig.4C, we used the average vein radius along with flow velocity to calculate key dimensionless flow metrics: Péclet (Pe) number (Fig.4D) (ratio of advection to diffusive transport), pulse frequency (Fig.4E) (i.e., pulsatility of flows), Reynolds (Re) number (Fig.4F) (ratio of inertial to viscous flows), and Womersley (Wo) number (Fig.4G) (ratio of pulsatility with respect to viscosity effects). Pe is similar between wing regions (Fig.4D), with the exception of the trailing edge of the forewing, where it is slightly higher. Pulse frequency, measured as the number of velocity peaks over time, is higher in regions of the wing where flow is returning back to the body, such as the trailing edge region (Fig.4E). Similarly, Reynolds number jumps by an order of magnitude, up to 0.04 (Fig.4E), in the trailing edge region of the forewings, where hemolymph is being pumped back into the body. While this flow is still dominated by viscous effects, it is less so in this region compared to the rest of the wing, underscoring the importance of the pumping organs in driving wing circulation. Wo is similar in all wing regions, except for a noticeable decrease in the wing tip, where flow tends to slow down (Fig.5G). Hemolymph flow within both wings has a similar Wo to arterioles and venules in the human body.^35^

**Figure 5.**
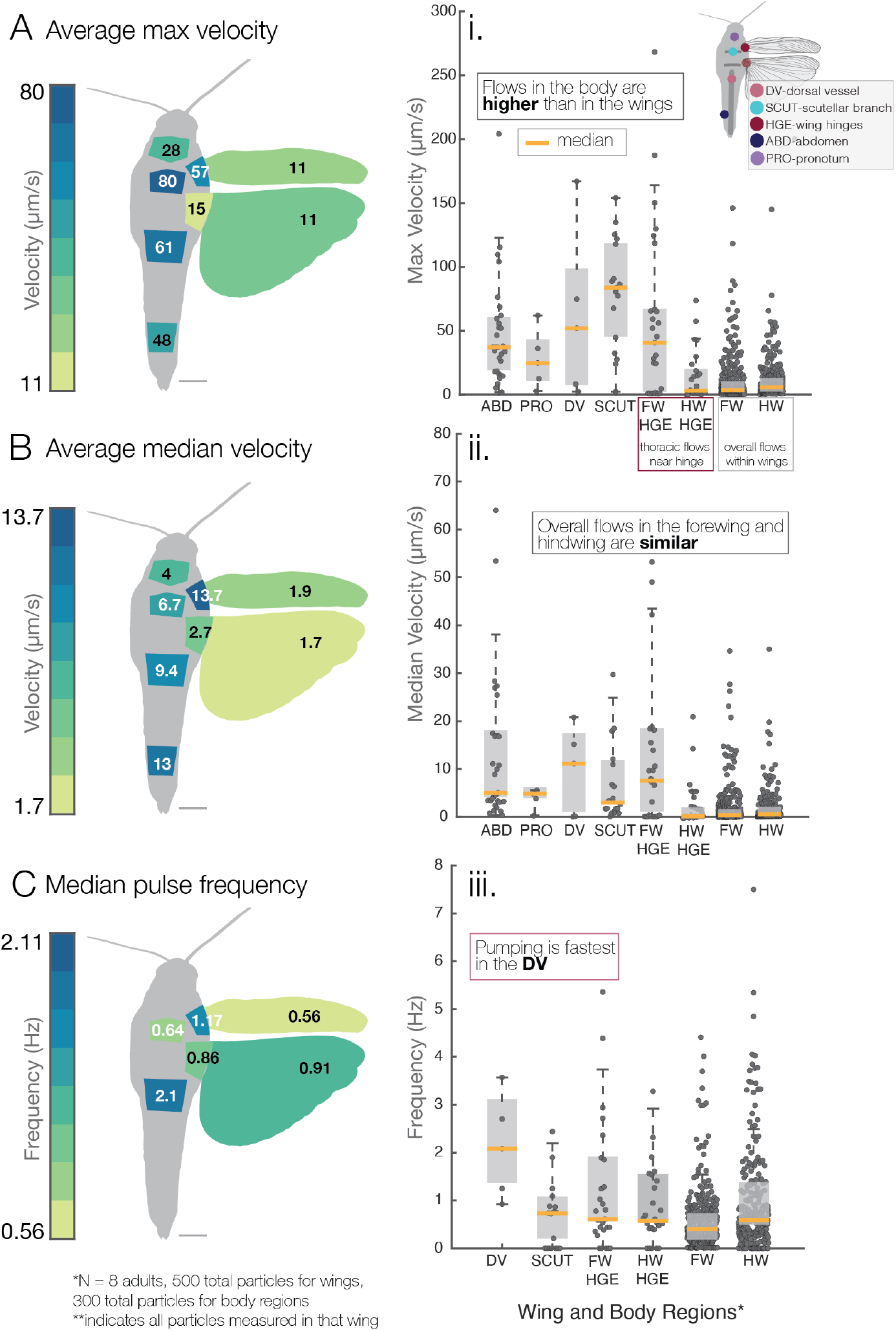
Flow metrics in the body versus wings. **(A)** Average of the peak particle velocities across the body and wings as a whole. A,i shows box and whisker plots (median in red) of particle data. Flows are faster within the body than in wings. **(B)** Average of the median velocities indicate faster flows in the body, but in B,ii medians overlap except with the dorsal vessel (DV). **C** Average pulse frequency (as calculated in Fig.4E) describes pumping is highest in the dorsal vessel (2.1 Hz), while the scutellar branch (SCUT), the return conduit for flow, and wing areas have similar frequencies. FW-THX indicates thoracic flows near the forewing base. Scale bar - 5 mm.

### The body and wing pumps display faster hemolymph flow than the wings

Hemolymph is pulled from the wing through the scutellar branches, and pumped back into the body by the thoracic wing hearts (Supp Movies 7-8). Flow enters the wing through thoracic openings that connect to the leading edge veins, and flow within the wings is constrained by the dense network of small veins. Thus it is not surprising that flow velocities measured within the thorax and the rest of the body were much faster than those in the wings (Figure 5). Flows in the thorax near the wing hinge were also turbulent, with mixing of in-going hemolymph with hemolymph in the thoracic cavity. The dorsal vessel and scutellar branches (Fig.2B, Supp. Movie 7)) both displayed significantly higher flow velocities (Fig.5A,B) than those measured in the forewing, hindwing, or hindwing hinge (paired t-test, P<0.001). Flow velocities within the abdomen were also higher than those within the forewing, hindwing and hindwing hinge, whereas flows in the pronotum were not statistically different from any other sampled regions except for the scutellar branch (P<0.05).

Due to the role that the pumping organs play in driving hemolymph flow, differences in pulse frequency (i.e., pulsatility) among sampled regions of the body and wings are similar to those seem in flow velocity (Fig.5C). We measured a mean pulse frequency of 2.1 Hz in the dorsal vessel, which is significantly higher than the pulse frequency measured in the return conduits from the wing (scutellar branches). This frequency is somewhat higher than previously measured dorsal vessel pumping frequencies (0.92 Hz)^36^, but measurements of pumping frequency may vary depending on the specific location sampled along the dorsal vessel. Pumping frequencies of the scutellar branch, wing hinges, and wings are not significantly different from each other (Fig.5C, paired t-test). The similarity in hemolymph pulsing among these regions is also confirmed by similar Wo values calculated for the fore- and hindwing regions near the hinges (Figure 4G).

## Discussion

This detailed, hemodynamic analysis of global and local flow behaviors within an insect wing is the first of its kind, and sets a foundation for future studies examining the diversity of hemolymph circulation strategies that other insects may display. Our results are consistent with Arnold’s 1960 work on different insect species, demonstrating an overall circuitous flow pattern within locust wings (Fig. 3C), with hemolymph actively pumped out of the trailing edge veins and passively flowing into the leading edge veins. However, we also show that on a local level, hemolymph circulates through every vein within the wing, even the smallest cross-veins, and we identify three different types of local flow behaviors: pulsatile, continuous, and leaky flow. Despite the straightforward pattern of circuitous flow throughout the whole wing, local flow behaviors within individual veins are complex and time-varying, with several different flow behaviors present in different combinations within each region of the wing (Fig. 3D). Leakiness, a local flow behavior where hemolymph moves out of the leading edge veins and into membrane regions was described qualitatively in a brief note by Arnold in 1963^34^, but the dynamics of flow into and out of these pseudo-stigmas in insect wings is quantifed for the first time here. These “false sinuses” are thought to be regions of potential aerodynamic importance, where additional mass in the leading edge acts as an “inertial regulator” of wing pitch during flapping flight.^8, 34^ Leakiness is not constrained to pseudo-stigmas, and may also be present in the leathery tegmens or elytra (i.e., modified forewings of beetles), where tubular veins are absent from much of the wing.^11^ In contrast, dragonflies display a “true” sinus in the form of the pterostigma, a thickened, rectangular piece of cuticle near the leading edge of the wing, which forms a sinus where hemolymph pools.

Given the fact that most wing veins are not symmetric tubes, the Péclet, Reynolds, and Womersley numbers calculated here may not represent holistic flow characteristics. Future studies incorporating high-resolution x-ray tomography to visualize internal vein tissues in unprecedented detail^7^ would allow for accurate physical measurements of vein structure that could be used to more precisely evaluate the viscous flow regimes that characterize flow throughout the wings.

### Future: Environmental pressures and scaling of living wing networks

In many insects, wings become brittle with age and some wing regions appear to be devoid of hemolymph.^37^ Insects also display cumulative, non-repairable wing damage due to collisions, with pollinators that frequently fly through vegetation losing progressively larger amounts of wing area over time^38^ Wing damage, and the hemolymph clotting and desiccation that result, can lead to changes in the material properties of insect wings, with wing stiffness increasingly dramatically after wings are dessicated for as little as five minutes^39^; however, the effects of wing dessication on the internal, physiologial processes within wings remain unknown. Wing hydration also depends strongly on the waxy outer layer of the wing cuticle, which could be affected by pesticides and other environmental contaminants, and the weakening of this protective layer would lead to desiccation and changes in wing mechanics and physiology.

In addition to the effects of environmentally-mediated dessication on wing mechanical properties, hemolymph flow dynamics and the living systems that depend upon this flow could be affected by varying environmental temperatures. For example, consider a pollinator like the crepuscular hawkmoth that feeds at dawn and dusk, or a bumblebee that forages from early morning through evening in frigid, high-altitude meadows. A variety of different adaptations have been described that allow pollinating insects to continue feeding in the cooler temperatures characteristic of crepuscular periods and inclement weather, including physiological features that allow for flight at low temperatures and in low humidity, and adaptations for sensing and navigating between flowers in dim light.^40^Do insects adapted to fly in these types of conditions experience temperature-dependent changes in hemolymph circulation, or do they display adaptations to maintain consistent, effective circulation despite varying environmental temperatures? Extending this further, do flying insects living in regions with extreme climates (high or low temperatures, humidity levels, etc.) face unique pressures on their wing systems that may require adaptive changes in flow dynamics?

Finally, little is known about how the wide differences in body size, extending over several orders of magnitude among insect species, and the equally wide variation in life history strategies and flight behaviors, may affect patterns of hemolymph flow within wings. For example, the dragonfly, a highly maneuverable aerial predator, may require relatively fast hemolymph flows to supply the extensive network of sensory structures throughout its wings.^6, 10^ In comparison, a migratory insect such as a monarch butterfly, which often glides along air currents and needs to maximize energetic efficiency to travel long distances without feeding, may instead benefit from slower wing hemolymph flows, which require less active pumping..In addition, flow speeds are likely to vary widely within insect orders such as Lepidoptera, which display enormous variation in body size. Hemolymph flow speeds and pulsatility are known to differ between large lepidopterans like the Atlas moth^24^ and smaller species such as the painted lady butterfly^4^, but additional, comparative studies are needed to understand how hemolymph flow dynamics scale with body size, and how venation patterns and internal vein structure affect this relationship.

An insect wing is essentially a soft-bodied microfluidic device, composed of thin membranes and tubes, that develops over time and changes shape dynamically - both during metamorphosis and in adulthood (particularly in species where the adult wings can fold). Insect wings are deployed during ecdysis with the wing venation networks intact, a process that could inspire new technologies in the burgeoning field of microfluidics.^41^ During metamorphosis, the adult wing becomes fully formed, but it remains folded into a complex, origami-like structure that must be unfurled hydraulically. This active process, which lasts for about 40-60 minutes in many insects, relies upon the network of tubular wing veins to pressurize the wing with hemolymph.^42^ Relatively little is known about the mechanics or hormonal triggers involved in this process (outside of *Drosophila*) ^1^, but the potential applications of an improved understanding of wing expansion extend from small, biomedical devices to gigantic, autonomously unfolding satellite solar panels.

Every flight behavior performed by winged insects, from predation to pollination, relies on functioning wings. In an age of massive declines in insect populations and diversity due to industrialization, climate change, and disease, additional investigations into the living networks within the complex, yet fragile insect wing will only benefit our understanding of the unique role that this structure plays, and the external pressures that may affect its ability to function properly.

## Methods

### Care of insects and fluorescent particle injection

*Schistocerca americana* nymphs (2nd - 4th instars) were maintained at 30 - 35°C (16:8 hr light cycle), obtained from USDA (Sydney, Montana), and reared in accordance with USDA Aphis permits (#:P526P-16-04590). Once nymphs eclosed, adults were placed in a separate enclosure. Adults were regularly fed romaine lettuce, which supplied both nutrition and water.

To prepare for fluorescent microscopy, adult *S. americana* were briefly anesthetized with carbon dioxide and placed ventral side up. A small hole was drilled (approx. 0.096 mm^2^) in the second or third abdominal segment using an insect pin to accommodate injection. 6-10 microliters of a mixture of green fluorescent particles (Thermo Scientific, 1.05 g/cm^3^, fluorescence 589 nm) were injected using a 2.5 *μ*m Hamilton glass syringe with a pulled borosilicate capillary tube (Fig.2A). This mixture contained neutrally buoyant polystyrene particles of sizes 3 *μ*m and 6 *μ*m.

The mixture allowed flow to be observed at both large and small focal distances (Fig.2). After injection, *S. americana* were quickly restrained with modeling clay, allowing the live insect to remain at rest without moving or causing harm to itself. Fore and hind wings were spread and sandwiched between two glass slides (7.5 cm by 5 cm), at approximately a planar position, to simulate a wing-extended flight posture and allow for visualization of hemolymph flow (Fig.2). Due to an insect’s open circulatory system (Fig.1), injected particles flowed readily with hemolymph and were observed moving in and out of pumping organs, the body, and appendages.^23^

### Filming particle across insect wings

Particle movement was captured in 8 adult *S. americana* (approx. 3-5 months old) on a Zeiss AxioZoom Fluorescent Microscope at the Harvard Center for Biological Imaging (Cambridge, MA). Due to focal constraints, particle sizes, and clarity of the venation, no more than a third of the wing could be viewed at a time (approx. 300 mm^2^ for the hindwing and 10-50 mm^2^ for the forewing). Thus, movies were captured in a tiled fashion (wing base to wing tip) across the span and chord of the wing (Fig.2C). In practice, one can trace and follow a particle from wing hinge to the wing tip and back again; however visibility, frame rate, and file saving time constrained tracking distance within the wing. Frame rate ranged from 10-100 frames per second, where higher frame rates were necessary to capture rapid flow at the wing hearts and leading edge veins.

Approximately 800 particles from 228 movies of 8 individual adult grasshoppers were tracked and quantified for position, velocity, and acceleration. Particles were autotracked when possible using DLTdv5, a MATLAB based point-tracking program.^33^ Maximum velocity, path sinuosity (a ratio of distances), and net rate of fluid movement were calculated. Particle trajectories were measured individually, placed on a normalized wing coordinate system, and categorized into five wing regions (Fig.3B,C) and major body regions (Fig.2C). Velocity data were smoothed using a moving-mean function (MATLAB) with window-length of 5.

### Calculating wing region metrics

Across the five wing regions (Fig. 4), velocity (peak and median), average vein radii, pulse frequency, Péclet number, Reynolds number, Pulse frequency, and Womersley number were calculated. Velocity (*v_max_, μ*/s): how fast particles move through a region (maximum and median velocities were calculated to show range of particle movement). Vein radius was determined by taking an average of 25 vein diameters within a wing region. Pulse frequency (*f*, Hz) measures flow pulsatility, where the number of peaks in a velocity trace (over time) were used as an indication of pulsatility (Fig.4E. Velocity traces were normalized by *v_peak_* and peaks were detected using a threshold value of 0.3, which captured most apparent pulsatility. Péclet number reflects the ratio of viscous flows to diffusive transport. Reynolds number is a ratio of inertial to viscous fluid forces. Womersley detects the relevance of pulsatility to the viscous effects in a flow. The equations used are as follows:

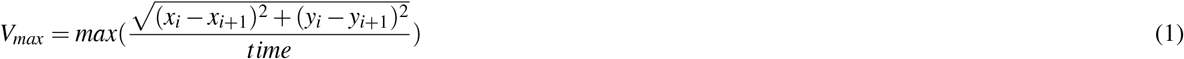

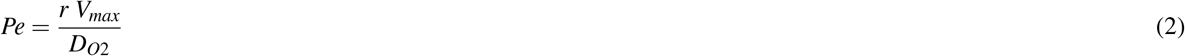

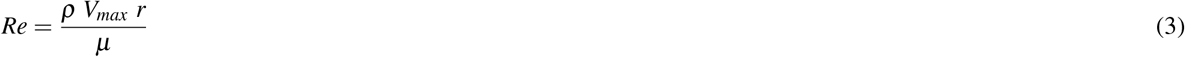

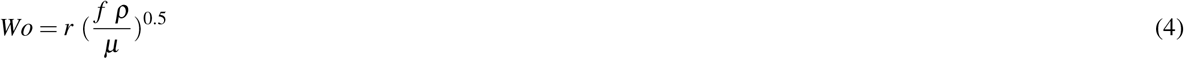

where points (*x_i_,yi*) and (*x*_*i*+1_,*yi*+1) are trajectory points through which a given particle travels, *ρ* is the density of water, *μ* is the dynamic viscosity of water, *r* is the average radii per wing region, *f* is the average pulse frequency per wing region, and *D*_*O*2_ is the diffusion coefficient of oxygen into water.

### Using cell tracking algorithms to capture particle movement

A detailed methodology on multi-parametric particle tracking algorithms can be found in our previous work^43–45^. Background subtraction was first applied to each frame in the time series to address low contrast ratios (CR) and compensate for uneven spatial illumination levels. A frame-wise linear intensity adjustment was applied, such that 1% of the total pixels were saturated, accounting for temporal fluorescence decay due to photobleaching. A local Hessian matrix of the intensity was calculated for each pixel, and the particles were marked by negative *λ*2 values in the Hessian eigenmaps. A dynamic erosion procedure with an adaptive threshold was used to identify each intensity peak of all particles that were analyzed. Subsequently, a dilation procedure was used to expand the boundaries from the identified peaks until it captured the course boundary of each particle. Finally, the coarse segmentation was mapped back to the original resolution and refined. The refining expansion stopped either when the pixel intensity fell below 25% of the peak intensity within the particle, or when it met the edges detected by a Canny filter^46^. Our algorithm identifies the most probable correspondence between particles by taking into consideration the characteristics of each particle (brightness, area, diameter, and orientation.) in addition to the classic nearest-neighbor criterion as tracking parameters.

## Acknowledgements

This research was funded by the US National Science Foundation (NSF) through a NSF Graduate Research Fellowship and a NSF Postdoctoral Research Fellowship in Biology (NSF Award ID: 1812215). I would like to thank Dr. Stefan Jaronski (USDA) for a continuous grasshopper supply, thoughtful discussions, and support. Thank you to Dr. Missy Holbrook for recommending I find an “insect workhorse.” The authors would like to thank the entire lab for giving space to the grasshopper colony for two years and allowing insect experiments in her plant lab. We thank the Harvard Center for Biological Imaging for infrastructure and support, specifically Dr. Doug Richardson for his advice and time training. We thank Dr. Siddarth Srinivasan for his expertise in measuring biological flows and his time spent in helping train on the microscope. Thank you to Dr. Naomi Pierce for her advice and edits. We thank Dr. Jake Socha for his edits and advice. Lastly, especial thanks to Dr. Jacob Peters for his advice and support throughout the project.

## Author contributions

MKS conceived of research project, collected and analyzed data, and wrote manuscript. BJ analyzed data, added methods, helped to edit manuscript. PV contributed methods and manuscript edits. SAC and LM contributed significant edits and advising throughout project.

## Supplementary information

Videos showing hemolymph flow and particle movement within each wing region and the major pumping organs can be found in the Video Supplement.

